# Low-frequency variant functional architectures reveal strength of negative selection across coding and non-coding annotations

**DOI:** 10.1101/297572

**Authors:** Steven Gazal, Po-Ru Loh, Hilary K. Finucane, Andrea Ganna, Armin Schoech, Shamil Sunyaev, Alkes L. Price

## Abstract

Common variant heritability is known to be concentrated in variants within cell-type-specific non-coding functional annotations, with a limited role for common coding variants. However, little is known about the functional distribution of low-frequency variant heritability. Here, we partitioned the heritability of both low-frequency (0.5% ≤ MAF < 5%) and common (MAF ≥ 5%) variants in 40 UK Biobank traits (average N = 363K) across a broad set of coding and non-coding functional annotations, employing an extension of stratified LD score regression to low-frequency variants that produces robust results in simulations. We determined that non-synonymous coding variants explain 17±1% of low-frequency variant heritability 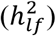 versus only 2.1±0.2% of common variant heritability 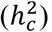, and that regions conserved in primates explain nearly half of 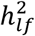 (43±2%). Other annotations previously linked to negative selection, including non-synonymous variants with high PolyPhen-2 scores, non-synonymous variants in genes under strong selection, and low-LD variants, were also significantly more enriched for 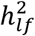 as compared to 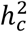. Cell-type-specific non-coding annotations that were significantly enriched for 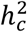 of corresponding traits tended to be similarly enriched for 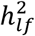 for most traits, but more enriched for brain-related annotations and traits. For example, H3K4me3 marks in brain DPFC explain 57±12% of 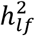 vs. 12±2% of 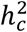 for neuroticism, implicating the action of negative selection on low-frequency variants affecting gene regulation in the brain. Forward simulations confirmed that the ratio of low-frequency variant enrichment vs. common variant enrichment primarily depends on the mean selection coefficient of causal variants in the annotation, and can be used to predict the effect size variance of causal rare variants (MAF < 0.5%) in the annotation, informing their prioritization in whole-genome sequencing studies. Our results provide a deeper understanding of low-frequency variant functional architectures and guidelines for the design of association studies targeting functional classes of low-frequency and rare variants.

## Introduction

Common variant (minor allele frequency (MAF) ≥ 5%) heritability is known to be concentrated into non-coding functional annotations that are active in relevant cell types or tissues (such as immune-cell-specific annotations in autoimmune diseases or brain-specific annotations in psychiatric diseases), with a limited role for common coding variants^1–9^. However, little is known about the functional architecture of low-frequency variants, which can have larger per-allele effect sizes when impacted by negative selection^10–16^. Recent studies have implicated large functional enrichments for low-frequency coding variants^17–20^, but their contribution to trait heritability is currently unknown; the contribution of low-frequency variants in cell-type-specific non-coding annotations is also unknown. Dissecting low-frequency variant functional architectures can shed light on the action of negative selection across functional annotations and inform the design of low-frequency and rare variant association studies^12,21^.

To investigate functional enrichments of low-frequency variants (defined here as 0.5% ≤ MAF < 5%), we extended stratified LD-score regression^5^ (S-LDSC) to partition the heritability of both low-frequency and common variants; our method produces robust (unbiased or slightly conservative) results in simulations. We applied our method to partition the heritability of low-frequency and common variants in 40 heritable traits from the UK Biobank^22–24^ (average *N* = 363K UK-ancestry samples, imputed using haplotypes from the Haplotype Reference Consortium^25^) across a broad set of coding and non-coding functional annotations^5,7,26–31^. We performed forward simulations to connect estimated low-frequency and common variant functional enrichments to the action of negative selection, and to predict the effect size variance of causal rare variants within each functional annotation.

## Results

### Overview of methods

S-LDSC^5,31^ is a method for partitioning the heritability causally explained by common variants across overlapping discrete or continuous annotations using genome-wide association study (GWAS) summary statistics for accurately imputed variants and a linkage disequilibrium (LD) reference panel. The method rests on the idea that if an annotation *a* is associated to increased heritability, LD to variants with large values of *a* will increase the *χ*^2^ statistic of a variant more than LD to variants with small values of *a*. Here, we extended S-LDSC to partition the heritability causally explained by low-frequency variants using GWAS summary statistics for accurately imputed and poorly imputed variants. We included separate annotations for low-frequency and common variants, and used WGS data from 3,567 UK10K samples^17^ as an LD reference panel to ensure accurate LD information for low-frequency variants in the UK-ancestry target samples analyzed in this study (see Online Methods).

More precisely, the expected *χ*^2^ statistic of variant *j* can be written as

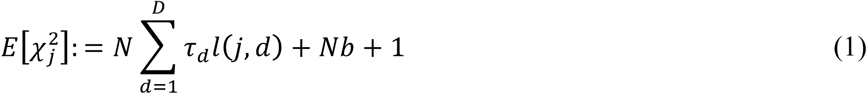

where *N* is the sample size of the GWAS study, *D* is the total number of annotations in the model (including both low-frequency and common variant annotations), *τ*_*d*_ is the effect size of annotation *d* on per-variant heritability (jointly modeled with all other annotations), 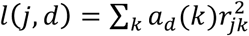 is the LD score of variant *j* with respect to continuous values *a*_*d*_(*k*) of annotation *d*, *r*_*jk*_ is the correlation between variant *j* and *k* estimated from the LD reference panel (e.g. UK10K), and *b* is a term that quantifies the contribution of confounding biases^32^.

We included 34 main annotations from our “baseline-LD model”^5,31^ (27 overlapping binary and 7 continuous annotations, including LD-related annotations) and 6 new binary main annotations: synonymous, non-synonymous, phastCons^26^ conserved elements in vertebrates, mammals and primates, and flanking bivalent TSS/enhancers^7^. We also considered 100bp and 500bp windows around binary annotations, duplicated all annotations for low-frequency and common variants, respectively, and included 10 common variant MAF bins (as in the baseline-LD model), 5 low-frequency variant MAF bins and one annotation containing all variants. We refer to the resulting set of annotations as the “baseline-LF model” (163 total annotations; Table S1 and Table S2; see Online Methods). We estimated the heritability causally explained by all low-frequency variants 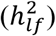 and the heritability causally explained by all common variants 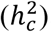; these computations did not include the heritability causally explained by rare variants (MAF < 0.5%). For the 33 main binary annotations, we computed their low-frequency variant enrichment (LFVE), defined as the proportion of 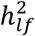 causally explained by variants in the annotation divided by the proportion of low-frequency variants that lie in the annotation, and common variant enrichment (CVE), defined analogously. We note that the choice of denominator in functional enrichment estimates merits careful consideration^33^ (see Online Methods), as comparisons of enrichment estimates with inconsistent denominators can produce incorrect conclusions^34^ (see Supplementary Note). Standard errors were computed using a block jackknife, as in our previous work^5,31^. Further details of the method are provided in the Online Methods section. We have released open-source software implementing the method, and have made our annotations publicly available (see URLs).

### Simulations to assess extension of S-LDSC to low-frequency variants

Although S-LDSC has previously been shown to produce robust results for partitioning common variant heritability using overlapping binary and continuous annotations^5,31,33^, we performed additional simulations to assess our extension to low-frequency variants. We first confirmed that S-LDSC with the UK10K LD reference panel produced unbiased heritability estimates for variants with MAF ≥ 0.5% in simulations using UK10K target samples, so that LD in the target samples and LD reference panel perfectly match (see Figure S1, Table S3, and Online Methods). We subsequently performed more realistic simulations using target samples from the UK Biobank interim release^22^ (*N* = 113,851 individuals with UK ancestry^35^, *M* = 1,023,655 variants from chromosome 1 with at least 5 minor alleles in UK10K), so that LD (and MAF) in the target samples and UK10K LD reference panel do not perfectly match. In detail, we (*i*) simulated quantitative phenotypes using integer-valued genotypes sampled from UK Biobank imputed dosages, (*ii*) computed summary statistics using UK Biobank imputed dosages (reflecting the impact of imputation uncertainty) and (*iii*) ran S-LDSC using our baseline-LF model and the UK10K LD reference panel (see Online Methods and Figure S2). S-LDSC was run either by restricting regression variants to accurately imputed variants (i.e. INFO score^36^ ≥ 0.99), as we recommended previously^5^, or by including all variants (regardless of INFO score). We focused our simulations on two representative annotations spanning roughly 1% of the genome: coding and ChromHMM/Segway weak-enhancer^37^ (enhancer), and considered various MAF-dependent architectures. Specifically, we simulated the variance of per-normalized genotype effect sizes proportional to (2*p*(1 − *p*))^1+*α*^ for a variant of minor allele frequency *p* (ref. ^38,39^), and allowed for different *α* values for variants inside and outside the coding/enhancer annotations; we conservatively specified our generative model to be different from the additive model assumed by S-LDSC (see Online Methods). For each of these two annotations, we simulated scenarios with no functional enrichment (“No Enrichment”) and scenarios with CVE roughly equal to 7x and lower LFVE (“Lower LFVE”), similar LFVE (“Same Enrichment”), or higher LFVE (“Higher LFVE”), respectively. For both coding and enhancer annotations, we observed that including all variants in the regression produced slightly conservative LFVE estimates and unbiased LFVE/CVE ratio estimates, while restricting to accurately imputed variants produced upward biases (Figure 1, Table S4); using other INFO score thresholds did not improve the results (Table S4). The slightly conservative 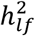 and LFVE estimates are due to LD-dependent architectures (coding and enhancer variants have lower than average levels of LD, as do other enriched functional annotations^31^), as we observed nearly unbiased estimates when creating shifted annotations with average levels of LD (see Online Methods; Figure S3). We thus recommend including all variants in the regression when running S-LDSC using the baseline-LF model. Our simulations indicate that this method is robust (unbiased or slightly conservative) in estimating low-frequency and common variant functional enrichments and LFVE/CVE ratios across a wide range of genetic architectures, even in the presence of poorly imputed variants, a target sample that does not exactly match the UK10K LD reference panel, and a MAF-dependent architecture that does not match the additive model assumed by S-LDSC; however, our estimates should not be viewed as rigorous lower bounds, as we have not simulated all possible genetic architectures.

**Figure 1:**
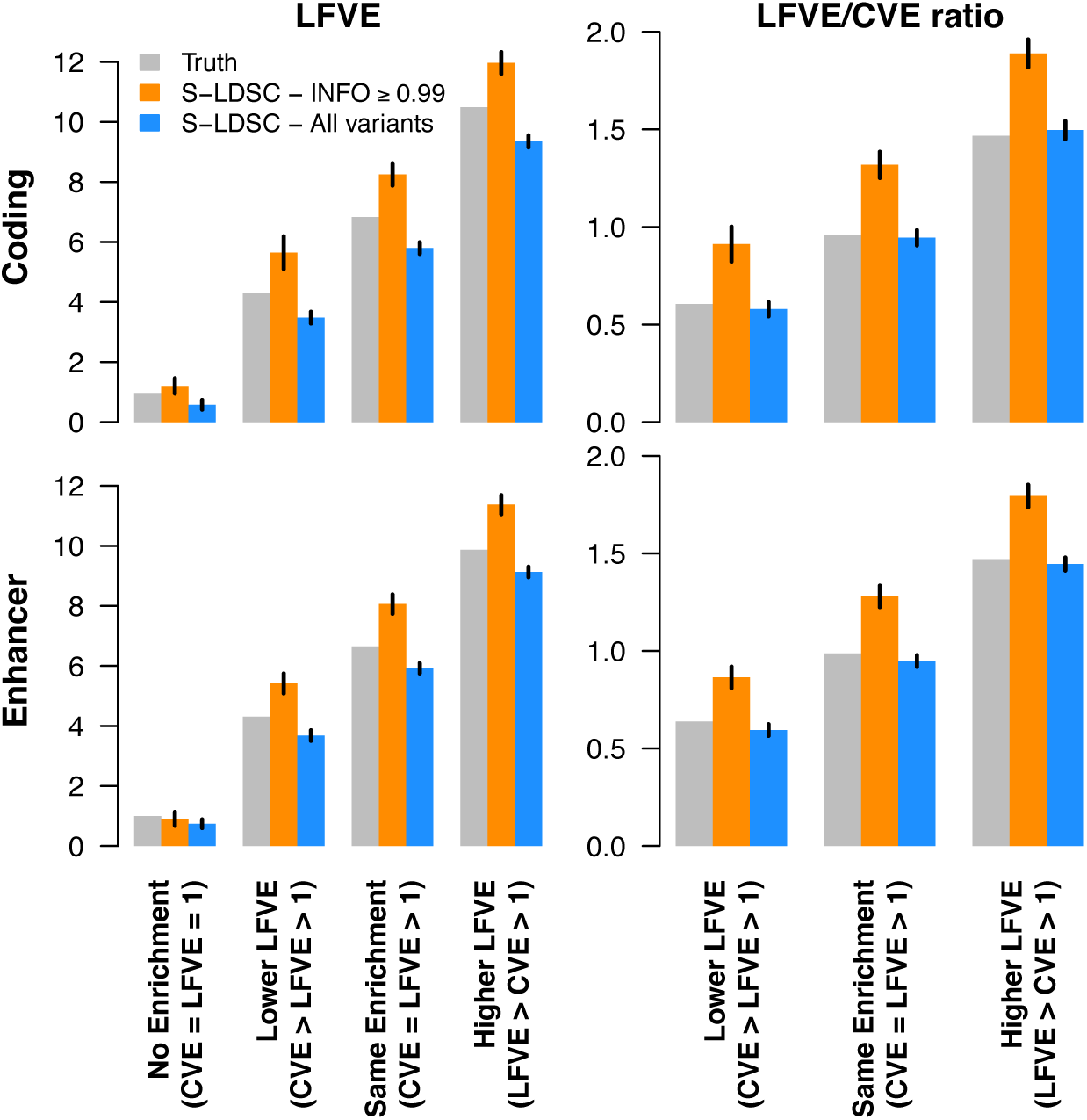
Simulations to assess low-frequency variant enrichment estimates. We report estimates of LFVE and LFVE/CVE ratio in simulations under a coding-enriched architecture (first row) or enhancer-enriched architecture (second row). We considered four different simulation scenarios (see main text). S-LDSC was run either by restricting regression variants to accurately imputed variants (S-LDSC – INFO ≥ 0.99), or by including all variants (S-LDSC – All variants). We do not report LFVE/CVE ratio for the No Enrichment simulation (CVE = LFVE = 1) due to unstable estimates; however, all analyses of real traits in this paper focus on annotations with significant CVE. Results are averaged across 1,000 simulations. Error bars represent 95% confidence intervals. Numerical results for 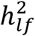, 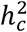, LFVE, CVE and LFVE/CVE ratio are reported in Table S4.

### Partitioning low-frequency and common variant heritability of UK Biobank traits

We applied S-LDSC with the baseline-LF model to 40 polygenic, heritable complex traits and diseases from the full UK Biobank release^23^ (average *N* = 363K; Table S5). Analyses were restricted to the set of 409K individuals with UK ancestry^23^ to ensure a close ancestry match with the UK10K LD reference panel, as LD patterns of low-frequency variants are expected to vary across European populations^40,41^. Summary statistics were computed by running BOLT-LMM v2.3 (ref. ^24^) on imputed dosages, and made publicly available (see URLs). S-LDSC results were meta-analyzed across 27 independent traits (average *N* = 355K; see Online Methods). We observed a roughly linear relationship between estimates of 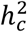 and 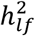 (Figure 2 and Table S5), with low-frequency variants explaining 6.3±0.2x less heritability and having 4.0±0.1x lower per-variant heritability than common variants on average. These ratios are consistent with a model in which the variance of per-normalized genotype effect sizes is proportional to (2*p*(1 − *p*))^1+*α*^ (ref. ^38,39^) with *α* = −0.37 (95% confidence interval [−0.40; −0.34]; similar to previous *α* estimates from raw genotype-phenotype data^15,16^), and consistent with a model in which low-frequency variants have smaller per-variant heritability but larger per-allele effect sizes^15,16,31,38,39^ (Figure S4).

**Figure 2:**
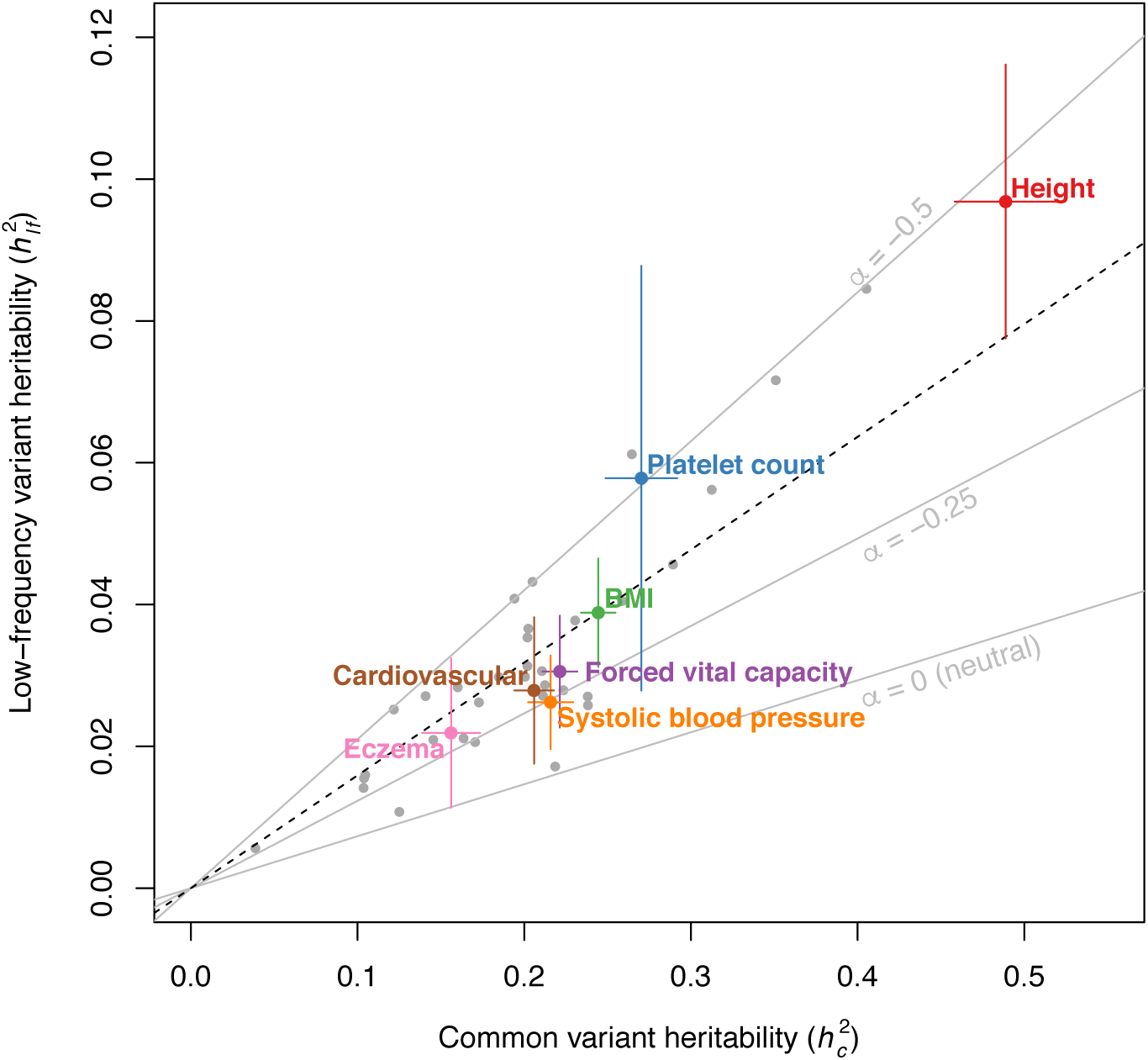
Common variant heritability 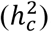 and low-frequency variant heritability 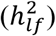 estimates for 40 UK Biobank traits. We report 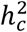 and 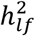 estimated by S-LDSC with the baseline-LF model for 40 UK Biobank traits (for binary traits, estimates are on the liability scale), with 7 representative independent traits highlighted. Error bars represent 95% confidence intervals. The dashed black line represents the ratio between 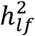 and 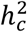 meta-analyzed across 27 independent traits (1/6.3). Grey lines represent expected ratios for different values of *α* (see main text). Error bars represent 95% confidence intervals. Numerical results are reported in Table S5.

We compared the LFVE and CVE of the 33 main binary functional annotations of the baseline-LF model, meta-analyzed across traits (Figure 3, Table S6). LFVE were highly correlated to CVE (*r* = 0.79) and larger than CVE on average (regression slope = 1.85). We identified 9 main functional annotations with significantly different LFVE and CVE (Figure 3, Table S6). Non-synonymous variants had the largest LFVE and largest difference vs.

**Figure 3:**
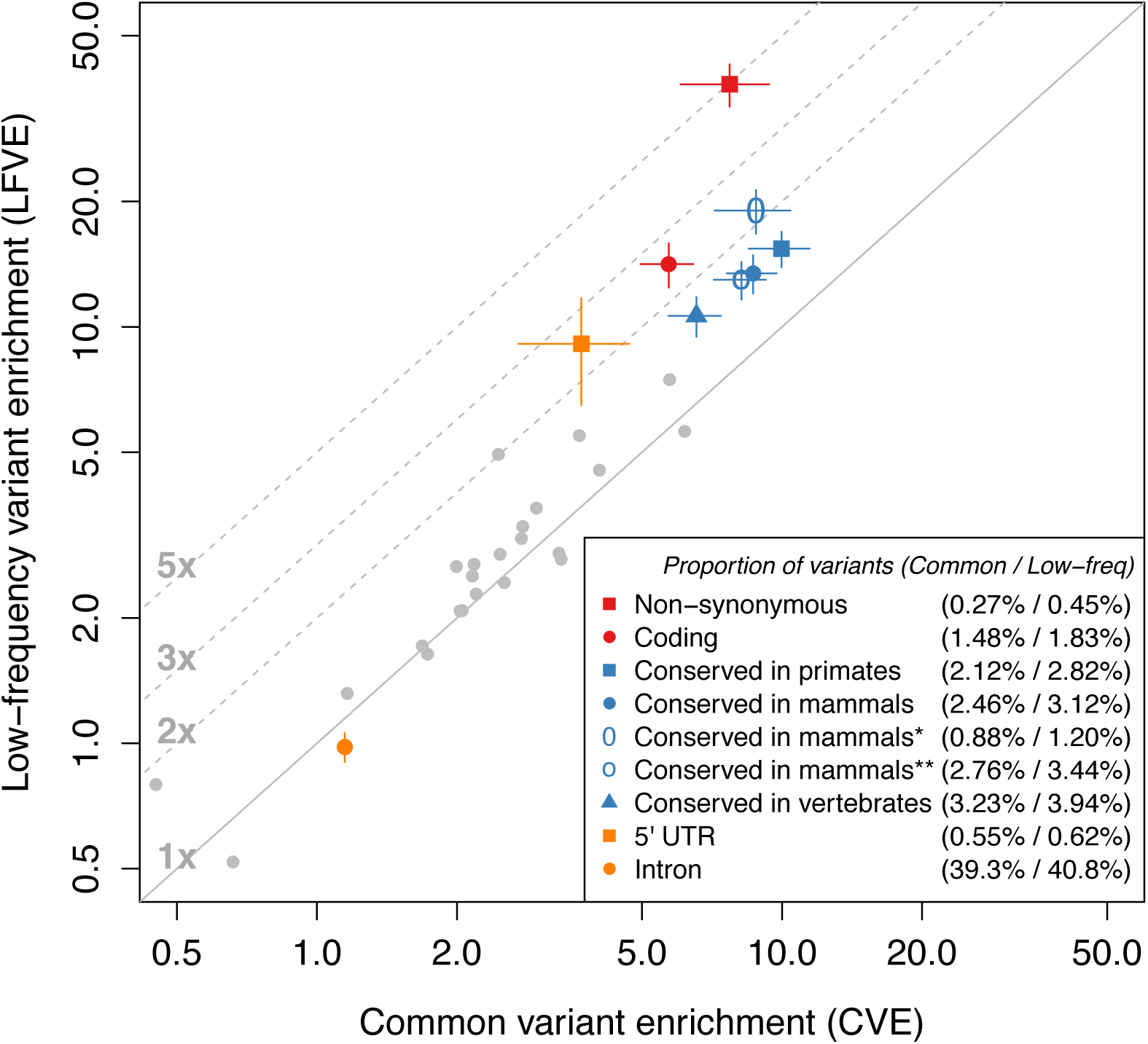
Functional low-frequency and common variant architectures across 27 independent UK Biobank traits. We plot LFVE vs. CVE (log scale) for the 33 main functional annotations of the baseline-LF model (meta-analyzed across the 27 independent traits), highlighting annotations for which LFVE is significantly different from CVE. Numbers in the legend represent the proportion of common / low-frequency variants inside the annotation, respectively. The first three conserved annotations are based on phastCons elements^26^, Conserved in mammals* is based on GERP RS scores^28^ (≥4), and Conserved in mammals** is based on Lindblad-Toh et al.^41^. The promoter flanking annotation has (non-significantly) negative LFVE and is not displayed for visualization purposes. The solid line represents LFVE = CVE; dashed lines represent LFVE = constant multiples of CVE. Error bars represent 95% confidence intervals. Numerical results are reported in Table S6.

CVE (5.0x ratio; LFVE = 38.2±2.3x, vs. CVE = 7.7±0.9x; *P* = 3 × 10^−36^ for difference). As non-synonymous variants comprise 0.45% of low-frequency variants vs. only 0.27% of common variants due to strong negative selection on non-synonymous mutations^42–44^ (see below), this difference is even larger when comparing the proportion of heritability they explain (8.2x ratio; 17.3±1.0% of 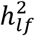, vs. 2.1±0.2% of 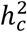; *P* = 5 × 10^−47^). Non-synonymous variants predicted to be deleterious by PolyPhen-2 (ref. ^27^) had larger LFVE and LFVE/CVE ratio than non-synonymous variants predicted to be benign (Figure S5).

We also observed LFVE significantly larger than CVE for coding variants (2.5x ratio; *P* = 1 × 10^−18^), 5’ UTR (2.5x ratio; *P* = 1 × 10^−4^) and the five main conserved annotations (three based on phastCons elements^26^, one based on a GERP RS scores^28^ ≥ 4, and one based on Lindblad-Toh et al.^29^; ratios 1.5x-2.2x; each *P* < 5 × 10^−7^; Figure 3, Table S6). Surprisingly, phastCons regions conserved in primates^26^ were more enriched than phastCons regions conserved in vertebrates or conserved in mammals^26^ (even though regions conserved in more distant species may be viewed as more biologically critical); the primates annotation explained nearly half of 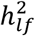 (2.8% of variants explaining 43.5±2.2% of 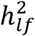, LFVE = 15.4x vs. 2.1% of variants explaining 21.2±1.6% of 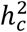, CVE = 10.0x; *P* = 1 × 10^−16^). We observed that the significantly larger LFVE (compared to CVE) for all 5 conserved annotations is mainly due to conserved regions that are coding, and that coding enrichments are similar for regions conserved across different species (Figure S6). On the other hand, the significantly larger LFVE and CVE for regions conserved in primates (compared to more distant species) is entirely due to conserved regions that are non-coding (Figure S6), consistent with the observation that many human enhancers have evolved recently^45–47^. Finally, we observed significantly smaller LFVE than CVE for intronic variants (0.85x ratio; *P* = 8 × 10^−5^). These results were generally consistent across the 40 UK Biobank traits analyzed: LFVE was larger than CVE for non-synonymous and regions conserved in primates in all 40 traits and for coding in 38 traits (non-significantly smaller for the other two traits), and LFVE was smaller than CVE for intronic variants in 33 traits (non-significantly larger for the other seven traits; Figure S7). We note that after removing the 9 annotations with significantly different LFVE and CVE, LFVE remained highly correlated to CVE (*r* = 0.83) and only slightly larger than CVE on average (regression slope = 1.10).

We also observed significantly larger enrichment/depletion for LFVE than for CVE in the first and/or last quintile of LD-related continuous annotations related to negative selection^31^ (Figure S8 and Table S7); our forward simulations from ref. ^31^ confirmed larger effects of low-frequency variants in these LD-related annotations (Table S8). Overall, our results suggest that LFVE is substantially larger than CVE only for annotations that are strongly constrained by negative selection, as the strongest differences were observed for coding and non-synonymous variants, which are known to be under strong negative selection^42–44^. A more detailed interpretation of the LFVE/CVE ratio is provided below (see Forward simulations).

### Cell-type-specific annotations dominate both low-frequency and common variant architectures

We sought to investigate the contribution to low-frequency variant architectures of cell-type-specific (CTS) annotations^1–9^ (i.e. reflecting regulatory activity in a given cell type) with excess contributions to common variant architectures. For each of the 40 UK Biobank traits, we selected the subset of 396 CTS Roadmap annotations^7^ with statistically significant common variant enrichment after conditioning on (non-CTS annotations in) the baseline-LD model^5,48^ (see Online Methods). We selected a total of 637 trait-annotation pairs, with at least one CTS annotation for 36 of 40 traits (25 of 27 independent traits) (Table S9); the 637 CTS annotations contained 2.7% of common variants and 3.0% of low-frequency variants on average (Table S10). We analyzed each of these trait-annotation pairs using the baseline-LF model (Figure 4a and Table S10). For the 25 trait-annotation pairs with the most statistically significant CVE for each of the 25 independent traits (critical CTS annotations), LFVE and CVE were similar, with LFVE 1.12x (s.e. = 0.13x) larger than CVE on average (other definitions of critical CTS annotations produced similar conclusions; see Figure S9).

**Figure 4:**
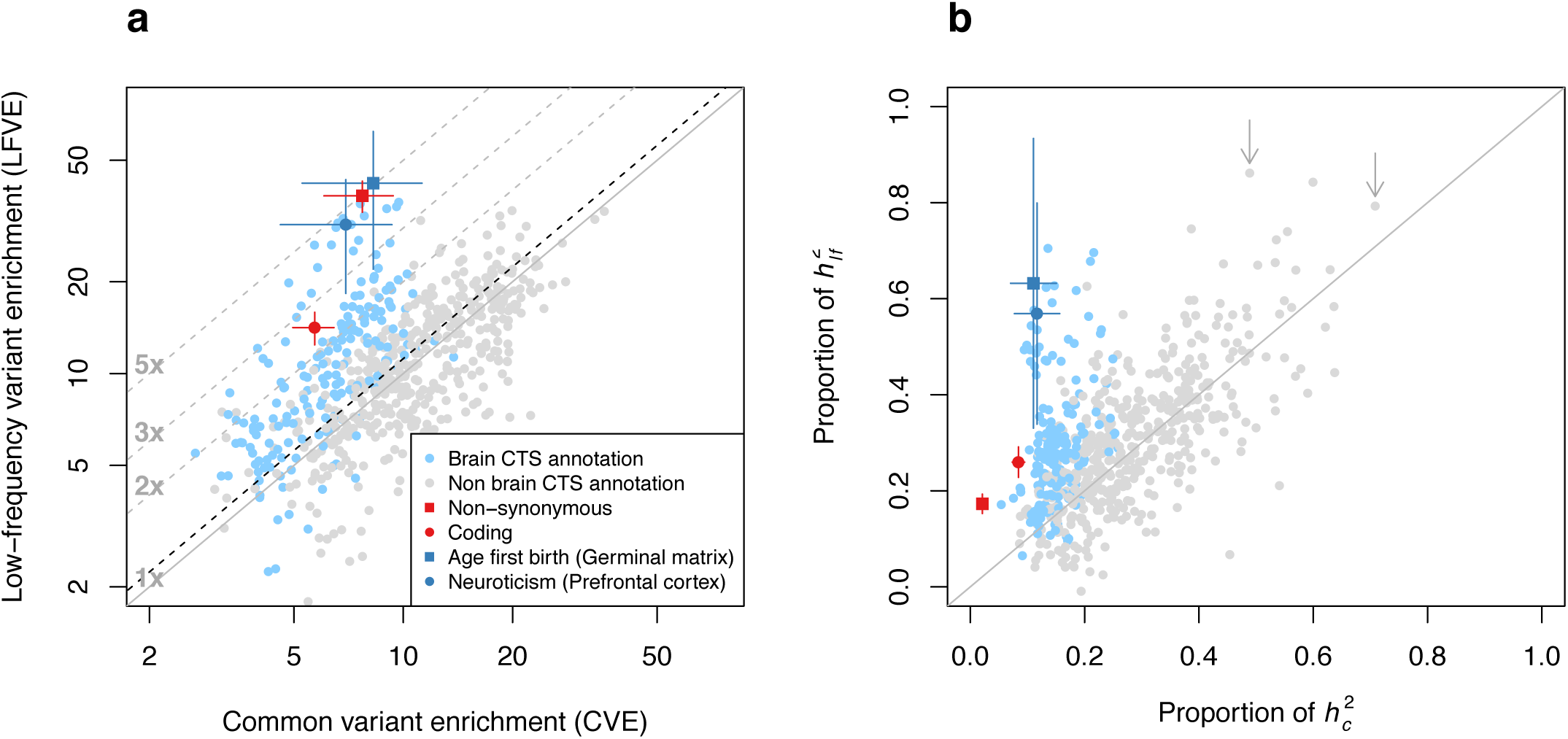
Low-frequency and common variant architectures of cell-type-specific (CTS) annotations. For 637 trait-annotation pairs with statistically significant common variant enrichment, we report **(a)** LFVE vs. CVE (log scale) and **(b)** proportion of 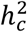 vs. proportion of 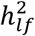 explained. The dashed black line in (a) represents the regression slope for 25 critical CTS annotations for independent traits (see main text). Brain-specific annotations are denoted in blue. Two trait-annotation pairs with LFVE significantly larger than CVE are denoted in dark blue (see main text); error bars represent 95% confidence intervals. The two arrows in (b) denote All autoimmune diseases (Regulatory T-cells) and Monocyte count (Primary monocytes) (see main text). Results for coding and non-synonymous annotations (meta-analysis across 27 independent traits) are denoted in red; error bars represent 95% confidence intervals. Numerical results are reported in Table S10.

We observed Bonferroni-significant differences (after correcting each trait for 1-53 annotations tested) for two traits, neuroticism and age of first birth in females. The most significant trait-annotation pairs were neuroticism and H3K4me3 in brain dorsolateral prefrontal cortex (4.4x ratio; LFVE = 30.8±6.4x, vs. CVE = 6.9±1.2x; *P* = 2 × 10^−4^ for difference; 56.9±11.7% of 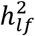, vs. 11.7±2.0% of 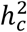), and age of first birth in females and H3K4me3 in brain germinal matrix (5.1x ratio; LFVE = 42.1±10.2x, vs. CVE = 8.3±1.5x; *P* = 0.001; 63.2±15.4% of 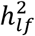, vs. 11.1±2.0% of 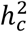). Interestingly, these two annotations (and 55 of all 62 CTS annotations with LFVE/CVE > 2) are brain-specific, implicating stronger selection against variants impacting gene regulation in brain tissues (see Forward simulations and Discussion).

While CTS annotations generally have only moderately large LFVE (e.g. smaller than non-synonymous variants; Figure 4a), they often explain a large proportion of 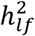 (e.g. larger than non-synonymous variants; Figure 4b) due to large annotation size, as with common variant enrichment. In particular, H3K4me1 in regulatory T-cells (3.7% of low-frequency variants) explains 86.2±20.8% of 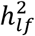 for All autoimmune diseases (vs. 3.4% of common variants explaining 48.9±9.1% of 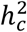), and H3K4me1 in primary monocytes (4.8% of low-frequency variants) explains 79.3±18.1% of 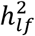 for monocyte count (vs. 4.6% of common variants explaining 70.8±8.6% of 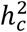; Figure 4b and Table S10). Thus, CTS annotations often dominate low-frequency architectures, analogous to common variant architectures^5,48^.

### Non-synonymous variants in genes under strong selection are highly enriched for heritability

Recent studies have identified gene sets that are depleted for non-synonymous variants^30,49–51^; these gene sets are enriched for ultra-rare non-synonymous mutations impacting complex traits^52–54^. To further investigate the connection between functional enrichment and negative selection, we stratified the CVE and LFVE of non-synonymous variants (Figure 3a) based on the strength of selection on the underlying genes. We considered 5 bins of estimated values of selection coefficients for heterozygous protein-truncating variants^30^ (*s*^*het*^), with 3,073 protein-coding genes per bin, and added annotations based on non-synonymous variants within each bin to the baseline-LF model (see Online Methods). We determined that both the LFVE and CVE of non-synonymous variants correlated strongly with the predicted strength of selection on the underlying genes (Figure 5 and Table S11). In particular, we observed extremely strong enrichments for non-synonymous variants in genes under the strongest selection (bin 1: LFVE = 102.0±7.9x and CVE = 41.5±4.8x). However, the LFCE/CVE ratio was smaller for non-synonymous variants in genes under the strongest selection (bin 1: 2.5x) than in genes under the weakest selection (bins 4+5: 5.8x); we discuss this surprising result below (see Forward simulations). We obtained similar results when stratifying non-synonymous variants in genes under varying levels of selective constraint based on other related criteria^49–51^ (Figure S10).

**Figure 5:**
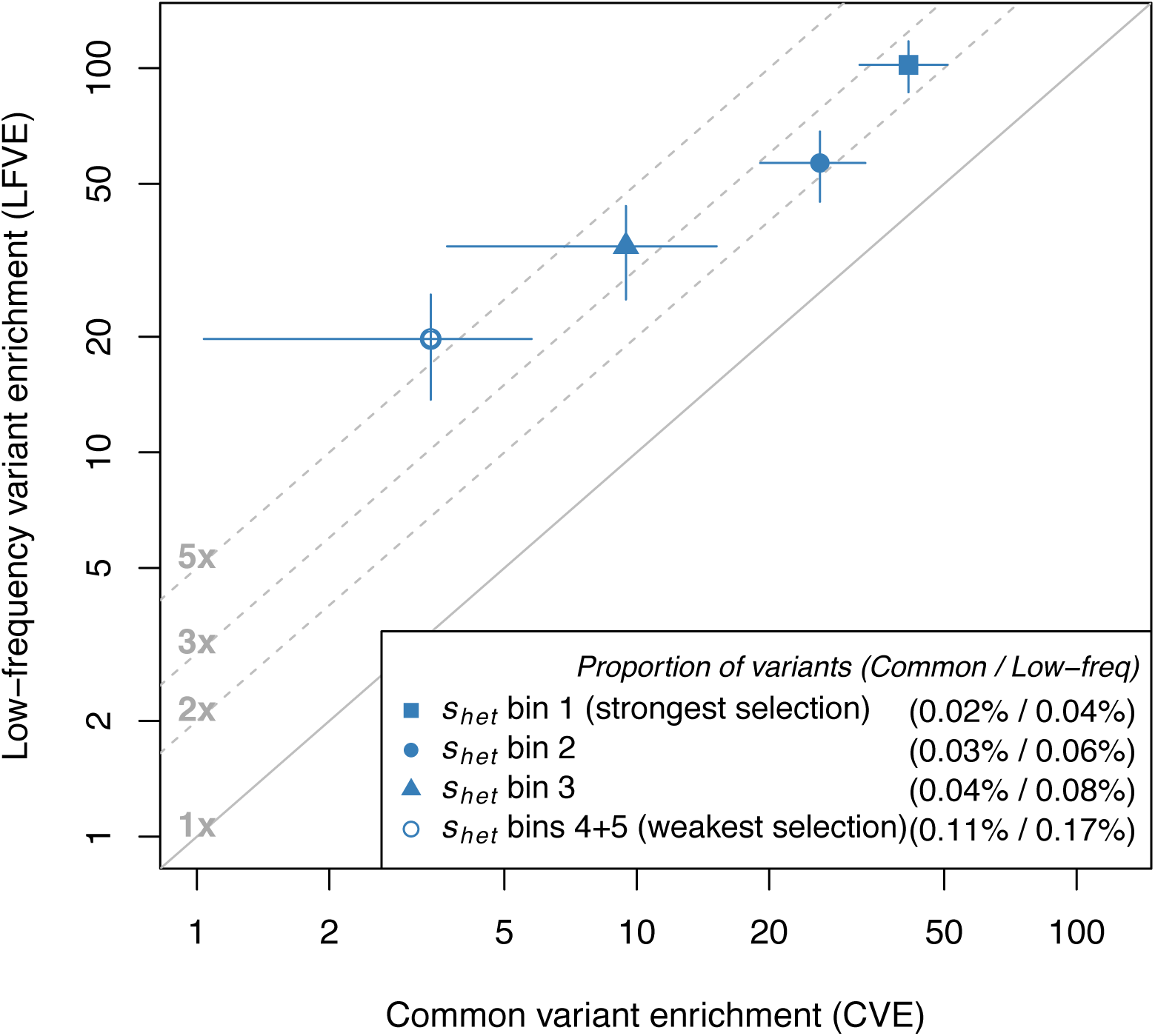
Low-frequency and common variant enrichments for non-synonymous variants vary with the strength of selection on the underlying genes. We report LFVE vs. CVE (log scale) for non-synonymous variants in 5 bins of *s*^*het*^ (see main text), meta-analyzed across 27 independent UK Biobank traits; bins 4+5 are merged for visualization purposes. Numbers in the legend represent the proportion of common / low-frequency variants inside the annotation, respectively. The solid line represents LFVE = CVE; dashed lines represent LFVE = constant multiples of CVE. Error bars represent 95% confidence intervals. Numerical results for each bin are reported in Table S11.

### Forward simulations provide interpretation of low-frequency variant enrichments

We hypothesized that the LFVE and CVE of different functional annotations would be informative for the action of negative selection, which constrains strongly selected variants to lower frequency^10,12,15,16,55–58^. To investigate this, we performed forward simulations^59^ using an out-of-Africa demographic model^60^ and a genetic architecture involving annotations mimicking non-synonymous variants (1% of the simulated genome), functional non-coding variants (1%), and ordinary non-coding variants (98%), with different respective distributions of selection coefficients *s* (Figure S11). For each of these three annotations we specified the probability for a *de novo* variant to be deleterious (π_*del*_), the mean selection coefficient for *de novo* deleterious variants 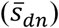 and the probability for a deleterious variant to be causal for the trait (π_*del:causal*_); the probability for a *de novo* variant to be causal for the trait is π = π_*del*_·π_*del:causal*_. Per-allele trait effect sizes were specified to be proportional to |*s*|^*τ*_*EW*_^ where *τ*_*EW*_ parameterizes the coupling between selection coefficient and trait effect size in the Eyre-Walker model^10^, implying that only deleterious variants have nonzero effects (see Online Methods). We investigated how the LFVE and CVE of the functional non-coding annotation varied as a function of the values of 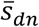 and π for that annotation. To achieve a realistic simulation framework, we fixed the remaining values of π_*del*_, 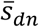 and π for the three annotations, as well as the value of *τ*_*EW*_, to values that we fit using our UK Biobank estimate of 4.0x larger per-variant heritability for common vs. low-frequency variants, as well as the LFVE and CVE of non-synonymous variants (38.2x and 7.7x, respectively). Specifically, we fixed π_*del*_ = 60% for the functional non-coding annotation (similar results for π_*del*_ = 40%; see Online Methods); π_*del*_ = 80% (ref. ^11^), 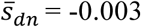 (ref. ^11^) and π = 8% for the non-synonymous annotation; π_*del*_ = 40%, 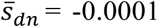 and π = 4% for the ordinary non-coding annotation; and *τ*_*EW*_ = 0.75. We note that our fitted value of *τ*_*EW*_ is larger than previous estimates^11,14,15,58^ (see Discussion).

We determined that the CVE of the functional non-coding annotation in our simulations depends on both 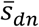 and π (Figure 6a), while the LFVE/CVE ratio depends primarily on 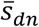 (Figure 6b). When *de novo* deleterious variants are under strong selection 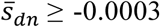, corresponding to LFVE/CVE ratio ≥ 1.2x; Figure 6b), the CVE depends primarily on π (Figure 6a), as the mean selection coefficient of deleterious common variants varies only weakly with 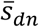 (since most deleterious common variants have 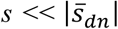; Figure 6c). Finally, we observed that functional non-coding annotations with similar CVE and LFVE tend to have causal variants with slightly stronger selection coefficients (i.e. 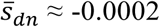) than ordinary non-coding causal variants 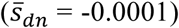, for which LFVE is lower than CVE (Figure 6b). We note that the LFVE/CVE ratio can be used to infer the mean selection coefficient of deleterious causal variants as a function of MAF (see Figure 6c), because this ratio depends primarily on 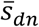 and because the selection coefficients of *de novo* deleterious causal variants are drawn from a distribution with mean 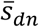.

**Figure 6:**
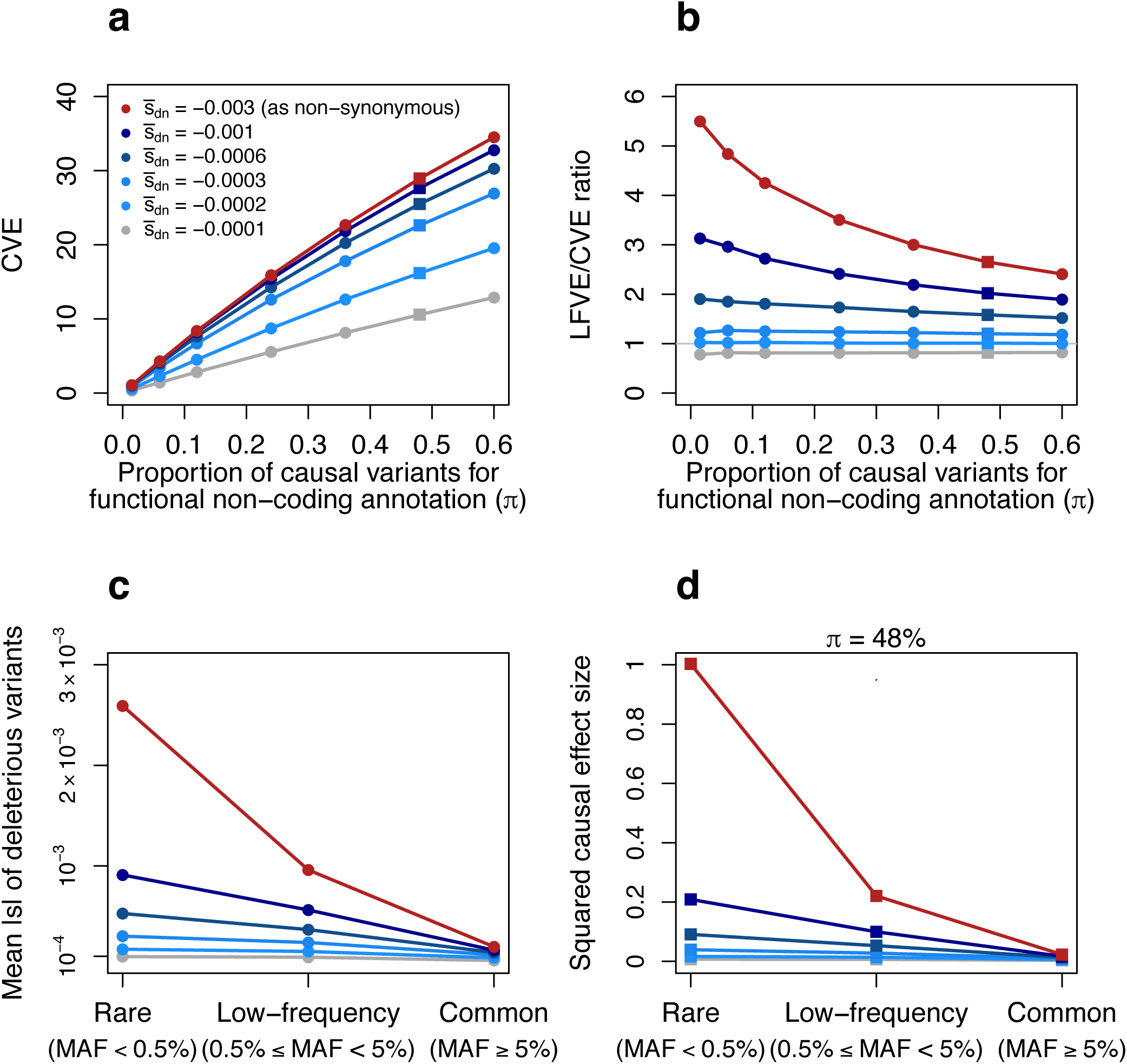
Forward simulations enable inferences about negative selection and rare variant architectures. Results are based on forward simulations involving an annotation mimicking functional non-coding variants, as well as other annotations (see text). **(a,b)** We report the CVE (a) and LFVE/CVE ratio (b) of the functional non-coding annotation as a function of the mean selection coefficient for *de novo* deleterious variants 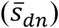 and the probability of a *de novo* variant to be causal (π) for this annotation. 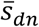 and π values for non-synonymous and ordinary non-coding annotations are described in the main text. **(c)** We report the mean absolute selection coefficient of deleterious variants in the functional non-coding annotation as a function of 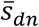 and MAF (rare, low-frequency, common). **(d)** We report the mean squared per-allele effect size of causal variants in the functional non-coding annotation (normalized by the mean squared per-allele effect size of rare causal non-synonymous variants) as a function of 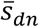 and MAF (rare, low-frequency and common). Red lines denote the value 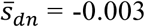 used to simulate non-synonymous variants, grey lines denote the value 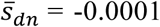 used to simulate ordinary non-coding variants (see main text). The value π = 48% used in (d) (see main text) is denoted via squares in (a) and (b). Numerical results are reported in Table S12.

Our forward simulations provide an interpretation of the LFVE/CVE ratios of different functional annotations that we estimated for UK Biobank traits and annotations. First, they confirm that non-synonymous variants (which are strongly deleterious^56^: large π_*de*l_ and 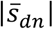) can have a limited contribution to common variant architectures (2.1% of 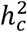) but a large contribution to low-frequency variant architectures (17.3% of 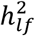) (Figure 3a). They are also consistent with stronger selection coefficients on 5’UTR variants and weaker selection coefficients on intronic variants (based on their LFVE/CVE ratios; Figure 3a). Second, they indicate that the proportion of causal variants (π) is larger for critical cell-type-specific (CTS) annotations than for non-synonymous variants (based on their CVE; Figure 4a), but that the causal variants in critical CTS annotations have only slightly larger selection coefficients than ordinary non-coding variants, except for some brain annotations that are under much stronger selection (much larger 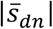, based on their LFVE/CVE ratios; Figure 4a). Third, they explain the extremely large CVE for non-synonymous variants inside genes predicted to be under strong negative selection^30^ (large *s*^*het*^; Figure 5), which are expected to correspond to genes with an extremely large proportion of deleterious non-synonymous variants (large π^del^, implying large π = π_*del*_·π_*del:causal*_). However, as noted above, this class of variants had a smaller LFVE/CVE ratio than that of non-synonymous variants inside genes predicted to be under weak selection (Figure 5), a surprising result that appears to suggest a smaller 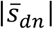 (Figure 6b) despite the extremely large value of π_*del*_. We performed additional forward simulations to show that a larger 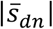 does not produce larger LFVE/CVE ratios for annotations with extremely large values of π_*del*_, due to more efficient background selection that decreases selection coefficients of low-frequency deleterious variants in such annotations (see Figure S12).

Although our focus is primarily on low-frequency variants (0.5% ≤ MAF < 5%), we also used our forward simulation framework to draw inferences about rare variant (MAF < 0.5%) architectures of non-coding functional annotations, based on LFVE and CVE estimates from UK Biobank (Figure 4a). Specifically, we compared the mean squared per-allele effect size of rare causal variants in annotations mimicking functional non-coding variants and non-synonymous variants, respectively. We focused on simulations with π = 48% for the functional non-coding annotation, because the CVE and LFVE/CVE ratios for the CTS annotations in Figure 4a (between 5 and 20, and between 1 and 2, respectively) roughly correspond to π = 48% and 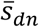 between 0.0002 and 0.0006 (Figure 6a-b). We inferred disproportionate causal effects of rare variants in annotations under very strong selection 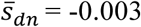, similar to non-synonymous variants^11^), with mean squared causal effect sizes 11x, 26x and 60x larger than annotations with 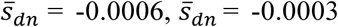 and 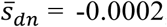, respectively (Figure 6d and Table S12; similar results for different choices of π, Figure S13). These results indicate that an annotation with large CVE needs to have even larger LFVE (e.g. LFVE/CVE ratio ≥ 2x, corresponding to 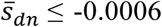 Figure 6b) in order to harbor rare causal variants with substantial mean squared effect sizes (e.g. only an order of magnitude smaller than rare causal non-synonymous variants; Figure 6d). Unfortunately, most of the non-brain CTS annotations that we analyzed do not achieve this ratio (Figure 4a), motivating further work on more precise non-coding annotations (see Discussion).

## Discussion

In this study, we partitioned the heritability of both low-frequency and common variants in 40 UK Biobank traits across numerous functional annotations, employing an extension of stratified LD score regression^5,31^ to low-frequency and common variants that produces robust (unbiased or slightly conservative) results. Meta-analyzing functional enrichments across 27 independent traits, we highlighted the critical impact of low-frequency non-synonymous variants (17.3% of 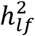, LFVE = 38.2x) compared to common non-synonymous variants (2.1% of 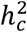, CVE = 7.7x); furthermore, regions conserved in primates^26^ explained nearly half of 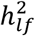 (43.5%, LFVE = 15.4x). Other annotations previously linked to negative selection, including non-synonymous variants with high PolyPhen-2 scores^27^, non-synonymous variants in genes under strong selection^30^, and LD-related annotations^31^, were also significantly more enriched for 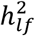 as compared to 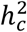. Finally, at the trait level, we observed that CTS annotations^7^,^48^ with significant CVE tend to have similar LFVE (or larger LFVE for brain-related annotations and traits; see below), implying that these annotations also dominate the low-frequency architecture.

We showed via forward simulations that the CVE of an annotation depends primarily on its proportion of causal variants (π), while its LFVE/CVE ratio depends primarily on the mean selection coefficient for *de novo* deleterious variants 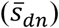, and thus to the mean selection coefficient of causal variants (Figure 6). These conclusions provide an interpretation of the enrichments that we estimated for UK Biobank traits and annotations. First, they confirm that non-synonymous variants can have a limited contribution to common variant architectures but a large contribution to low-frequency variant architectures; they also confirm higher selection coefficients on 5’UTR variants and lower selection coefficients on intronic variants. Second, they indicate that the proportion of causal variants is often larger for critical CTS annotations than for non-synonymous variants, but that causal variants in these annotations generally have only slightly larger selection coefficients than ordinary non-coding variants—with the exception of brain CTS annotations, whose larger LFVE (vs. CVE) implicates stronger selection on variants affecting gene regulation in the brain. This observation is consistent with the observation that brain enhancers interact with genes under stronger purifying selection^61^, and with the excess of rare *de novo* mutations in regulatory elements active in fetal brain in patients with neurodevelopmental disorders^62^. Third, they indicate that the extremely large CVE for non-synonymous variants inside genes predicted to be under strong negative selection^30^ is due to the high proportion of deleterious variants inside those genes, while their reduced LFVE/CVE ratio is due to the effect of background selection. Overall, our work quantifies the relationship between the strength of selection in specific functional annotations (both coding and non-coding) and low-frequency and common variant enrichment for human diseases and complex traits.

Our results on low-frequency variant functional architectures have several implications for downstream analyses. First, our results provide guidance for the design of association studies targeting low-frequency variants. Non-synonymous variants should be strongly prioritized at the low-frequency variant level^20^, as they explain a large proportion of 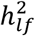 and directly implicate causal genes (and specifically implicate core disease genes rather than peripheral genes^63^), avoiding the challenge of mapping non-coding variants to genes^61,64–66^. However, we observed that all coding and UTRs variants jointly explained only 26.8±1.9% of 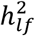 (Table S6), providing an indication of the proportion of low-frequency signal captured by whole-exome sequencing (WES) studies (we caution that this should not be viewed as a rigorous upper bound, as our estimates are slightly conservative in simulations; see Figure 1). This underscores the advantages of large GWAS (with imputed genotypes obtained using large reference panels such as HRC^25^), compared to WES or exome chip data, for querying low-frequency variation^14^. Furthermore, using functionally informed association tests that assign higher weight to low-frequency non-synonymous variants or CTS annotations should significantly improve power in these analyses^4,19,67^. Second, our results provide guidance for the design of association studies targeting rare (MAF < 0.5%) variants, which require large sequencing datasets^12^. While WES datasets have been successfully used to detect new coding variants, genes and gene sets associated to human diseases and complex traits^14,52,53,68,69^, there is an increasing focus on WGS that can capture rare non-coding variants^17,70–72^. However, our LFVE and CVE results for critical CTS annotations (Figure 4), coupled with our predictions of causal rare variant effect size variance (Figure 6d), suggest that in most instances these annotations do not harbor causal variants with large mean squared effect sizes (with brain-related annotations and traits as a notable exception; also see ref. ^62^), highlighting the need for more precise non-coding annotations for prioritization in WGS. As a first step towards this goal, we estimated the LFVE and CVE of annotations constructed using a wide range of recently developed non-coding variant prioritization scores^73–77^. We identified only one annotation, defined using the top 0.5% of Eigen scores^75^, with an LFVE/CVE ratio significantly larger than 1 (1.7x ratio; LFVE = 22.0±2.2x, vs. CVE = 13.0±1.4x; *P* = 7 × 10^−4^ for difference; Figure S14). However, even for this annotation, the LFVE/CVE ratio < 2 again implies that this annotation does not harbor causal variants with substantial mean squared effect sizes (only an order of magnitude smaller than rare causal non-synonymous variants; Figure 6d). Third, our results were consistent with strong coupling between selection coefficient and trait effect size (Eyre-Walker coupling parameter^10^ *τ*_*EW*_ = 0.75; robust to error bars in LFVE and CVE estimates, see Online Methods and Figure S15), implicating a larger impact of negative selection on complex traits than previously reported^11,14,15,58^ and much larger effect sizes for rare variants in functional annotations with strong selection coefficients. This can be explained by the fact that our inference procedure explicitly allows different distributions of selection coefficients for non-synonymous and non-coding variants 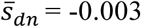 and 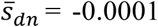, respectively; Figure S16). Finally, the different LFVE/CVE ratios that we inferred for different functional annotations suggest that it may be appropriate to allow annotation-specific *α* values when using the *α* model (per-normalized genotype effect size proportional to (*2p*(1 − *p*))^1+*α*^; ref. ^15,16,38,39^). In the extreme case of non-synonymous variants, we explored different choices of *α* values for non-synonymous and other variants, and determined that a value of *α* = −1.10 for non-synonymous variants and *α* = −0.30 for other variants provided the best fit to the UK Biobank estimated values of 4.0x larger per-variant heritability for common vs. low-frequency variants and LFVE = 38.2x and CVE = 7.7x of non-synonymous variants that we estimated in UK Biobank data (Table S13).

Although our work has provided insights on low-frequency variant architectures of human diseases and complex traits, it has several limitations. First, our simulations indicate that S-LDSC (using all variants) produces slightly conservative LFVE estimates for annotations with low levels of LD (Figure 1). However, this does not impact our main conclusions based on the LFVE/CVE ratio, for which S-LDSC produces unbiased estimates in our simulations (Figure 1). We further note that our S-LDSC model assumes a linear effect of each annotation and does not consider the effects of variants with MAF < 0.1% in the UK10K reference panel. Second, all of our analyses are based on UK-ancestry target samples and LD reference samples. Thus, more work is required to extend our approach to GWAS summary statistics from other European or non-European ancestries, together with available LD reference panels (such as the publicly available subset of HRC^25^ and TOPMed) that may not provide an exact match. Third, our method requires extremely large GWAS sample sizes with large matching LD reference panel, which are currently available only from the UK Biobank and UK10K data sets. Fourth, our method may sacrifice power by analyzing only GWAS summary statistics rather than the individual-level data that is readily available from UK Biobank. Although functional enrichment analyses of individual-level data can be performed using restricted maximum likelihood (REML) and its extensions^78–80^, those methods are applicable only to a small number of non-overlapping functional annotations; to our knowledge, all current methods that are applicable to a large number of overlapping functional annotations are based on summary statistics^81^, whereas analyzing one annotation at a time can produce severely biased results (see Figure 2b of ref. ^5^). Fifth, many of our analyses were meta-analyzed across 27 independent traits to maximize statistical power; this could mask trait-specific functional architectures, although results for non-CTS annotations were generally consistent across the 40 UK Biobank traits we analyzed (Figure S7). Sixth, the set traits that we analyzed includes only a limited number of diseases, due to the population-based design of UK Biobank (which limits power for diseases of lower prevalence); the analysis of larger case-control data sets would be of critical value, particularly for psychiatric disorders. Seventh, our inferences about negative selection and the effect size variance of causal rare variants in each annotation rely both on the Eyre-Walker model^10^ and a gamma distribution of selection coefficients^82^ (see Online Methods); large WGS datasets will be necessary to formally validate our extrapolation to rare variant architectures. Finally, our conclusions about CTS and non-coding annotations were limited by the availability and precision of regulatory annotations in relevant cell types. The generation of more comprehensive and more precise CTS annotations will aid our understanding of the non-coding genome and its role in human diseases and complex traits. Despite these limitations, our low-frequency and common variant enrichment results convincingly demonstrate and quantify the action of negative selection across coding and non-coding functional annotations.

## Online Methods

### Extension of S-LDSC to low-frequency variants

S-LDSC^5,31^ is a method for partitioning heritability explained by common variants across overlapping annotations (both binary and continuous^31^) using GWAS summary statistics. More precisely, S-LDSC models the vector of per normalized genotype effect size *β* as a mean-0 vector whose variance depends on *D* continuous-valued annotations *a*_1_,…, *a*_*D*_:

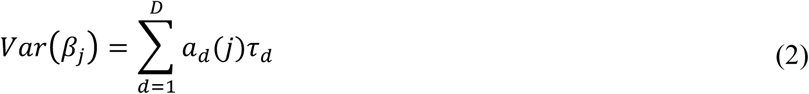

where *a*_*d*_(*j*) is the value of annotation *a*_*d*_ at variant *j*, and *τ*_*d*_ represents the per-variant contribution of one unit of the annotation *a*_*d*_ to heritability. We can thus perform a regression to infer the values of *τ* using the following relationship with the expected *χ*^*2*^ statistic of variant *j*:

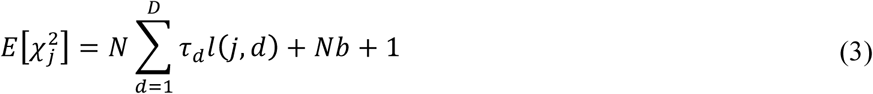

where 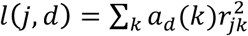 is the LD score of variant *j* with respect to continuous values *a*_*d*_(*k*) of annotation *a*_*d*_, *r*_*jk*_ is the correlation between variant *j* and *k* in an LD reference panel, *N* is the sample size of the GWAS study, and *b* is a term that measures the contribution of confounding biases^32^. Then, the heritability causally explained by a subset of variants *S* can be estimated as 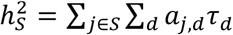. We note that this definition, used here to define and estimate 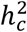 and 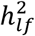, is different from the definition of “SNP-heritability” 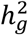 (ref. ^78^), which refers to the heritability tagged by a set of genotyped and/or imputed variants.

To allow different effects for low-frequency and common variants inside a functional annotation *a*_*d*_, we modeled the variance of the per normalized genotype effect sizes using different *τ*_*d*_ for these two categories of variants. In a case where we consider *D*_*f*_ functional annotations, we write:

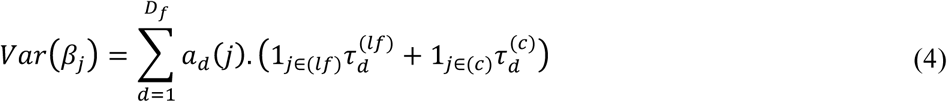

where 1_*j*∈(*lf*)_ (resp. 1_*j*∈(*c*)_) is an indicator function with value 1 if variant *j* is a low-frequency (resp. common) variant, and 0 otherwise, 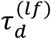 (resp. 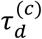) represents the per-variant contribution of one unit of the annotation *a*_*d*_ to the heritability explained by low-frequency (resp. common) variants. These parameters can be estimated using S-LDSC by writing equation (4) in the form:

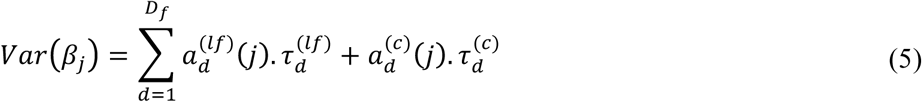

where 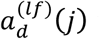 (resp. 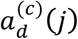) is an annotation equals to *a*_*d*_(*j*) if variant *j* is a low-frequency (resp. common) variant and 0 otherwise. In all analyses we also added one annotation containing all the variants, 5 MAF bins for low-frequency variants, and 10 MAF bins for common variants in order to take into account MAF-dependent effects^31,80,83^. For each functional binary annotation of interest *a*_*d*_, we compared its low-frequency variant enrichment (LFVE) and common variant enrichment (CVE), defined as the proportion of 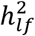 (resp. 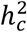) explained by the annotation, divided by the proportion of low-frequency (resp. common) variants that are in the annotation. Standard errors were computed using a block jackknife procedure^5^.

We emphasize that the choice of denominator (e.g. the proportion of low-frequency or common variants in an annotation) can have a large impact on the value of enrichment estimates, as demonstrated by two examples. First, coding regions are depleted for common variants (but less depleted for low-frequency variants), so that LFVE/CVE will be lower than (% of 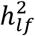)/(% of 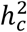); the same may be true for other annotations impacted by negative selection (see Results). A second example demonstrating that comparisons of enrichment estimates with inconsistent denominators can produce incorrect conclusions^34^ is discussed in the Supplementary Note.

Application of S-LDSC was performed using 3,567 unrelated individuals of UK10K data set^17^ (ALSPAC and TWINSUK cohorts) as an LD reference panel. This choice was made in order to ensure a close ancestry match between the target sample used to compute summary statistics (UK Biobank) and the LD reference panel (UK10K). We note that there is a difference between the LD reference panel (here UK10K, used to compute LD scores) and the imputation reference panel (here HRC, used to impute variants). Reference variants, used to estimate LD scores, were the set of 11,830,279 biallelic variants with MAF ≥ 0.1%. Regression variants, used to estimate annotation effect sizes *τ*, were the set of 8,523,464 UK Biobank variants with MAF ≥ 0.5% in UK10K. These MAF thresholds for reference variants and regression variants were assessed via simulations (see below). Variants with very large *χ*^*2*^ association statistics (larger than 0.001*N*), as well as variants in the major histocompatibility complex (MHC) region (chr6:25Mb-34Mb) were removed from all analyses. The main differences as compared to standard S-LDSC analyses on common variants are summarized Table S14.

### Baseline-LF model and functional annotations

We considered 34 main functional annotations from the baseline-LD model v1.1 (27 binary and 7 continuous annotations, including LD-related annotations; refs. ^5,31,84,85^), including coding, UTR, promoter and intronic regions, the histone marks monomethylation (H3K4me1) and trimethylation (H3K4me3) of histone H3 at lysine 4, acetylation of histone H3 at lysine 9 (H3K9ac) and two versions of acetylation of histone H3 at lysine 27 (H3K27ac), open chromatin as reflected by DNase I hypersensitivity sites (DHSs), combined chromHMM and Segway predictions (which make use of many Encyclopedia of DNA Elements (ENCODE) annotations to produce a single partition of the genome into seven underlying chromatin states), three different conserved annotations, two versions of super-enhancers, FANTOM5 enhancers, typical enhancers, and 6 LD-related continuous annotations (predicted MAF-adjusted allele age, level of LD in African populations, recombination rate, nucleotide diversity, a background selection statistic and CpG-content) (see Table S1).

In order to further dissect the set of coding variants, we annotated each coding variant using ANNOVAR^86^, and added one synonymous and one non-synonymous annotation (corresponding to “synonymous SNV” and “stopgain” or “stoploss” ANNOVAR outputs, respectively) into our model. (We note that the coding annotation in our baseline model comes from the UCSC coding track and includes UTR variants of protein coding genes, implying that some variants in this annotation do not belong to the synonymous or non-synonymous annotations). We also considered 4 new annotations based on phastCons^26^ conserved elements (46 way) in vertebrates, mammals and primates, and one annotation based on flanking bivalent TSS/enhancers from Roadmap data^7^ (see Table S15 for more information on the choice of annotations, as well as URLs). These 6 new annotations led to a total of 33 main binary annotations (see Table S1).

We included 500 bp windows around each binary annotation (except for annotations that are defined separately for each base pair) and 100 bp windows around four of the main annotations^5^, leading to a total of 74 main functional annotations. Then, all annotations were duplicated for low-frequency and common variants as described in equation (5), except for the predicted allele age annotation^87^ (which had too many missing values for low-frequency variants). Finally, we included one annotation containing all variants, 5 MAF bins of low-frequency variants and 10 MAF bins of common variants (see Table S16 for MAF bin boundaries). We thus obtained a set of 163 total annotations. We refer to this set of annotations as the “baseline-LF model” (see Table S2), which we used for all our S-LDSC analyses.

### Simulations using UK10K target samples to assess extension of S-LDSC to low-frequency variants

To obtain an initial assessment of the lowest MAF threshold for which S-LDSC with the UK10K LD reference panel might produce unbiased estimates, we first performed simple simulations using UK10K target samples (so that LD in the target samples and LD reference panel perfectly match) to investigate possible biases in 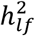 estimates when using different MAF thresholds to define low-frequency variants. We simulated quantitative phenotypes from chromosome 1 of UK10K data^17^ (3,567 individuals and 1,041,378 variants with allele counts greater or equal to 5, i.e. MAF ≥ 0.07%) by setting trait heritability to *h*^2^ = 0.5, selecting *M* = 100,000 causal variants, and simulating the variance of per-normalized genotype effect sizes proportional to *2p*(1 − *p*))^1+*α*^ for a variant of frequency *p*. We considered three values of *α*: −0.25 (close to the −0.28 value previously estimated^80^), −0.50 and −0.75 (with the latter values giving higher weight to low-frequency variants), and performed 5,000 simulations for each value of *α*.

To determine at which MAF thresholds we can accurately estimate heritability above the MAF threshold using UK10K as an LD reference panel, we constructed 6 different annotation models, with heritability variants (used to compute estimates of heritability and enrichment) and reference variants (used to compute LD scores) defined using MAF ≥ 0.1%, 0.2% or 0.5% (see Table S3 for the 6 different models). We estimated 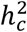 and 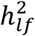 using S-LDSC and three different sets of regression variants (included in the regression, see equation (1)), also defined using MAF ≥ 0.1%, 0.2% or 0.5%. We note that our set of reference variants did not contain causal variants with a MAF between 0.07% and the selected MAF threshold, consistent with a realistic scenario in which causal rare variants may be absent from the reference data set. We determined that in these simulations, restricting to regression variants with MAF ≥ 0.5%, reference variants with MAF ≥ 0.1%, and heritability variants with MAF ≥ 0.5% provided unbiased estimates of 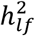 (Figure S1; we observed upward bias when considering MAF thresholds of 0.1% or 0.2% for heritability variants, Table S3). We thus used these MAF thresholds for further simulations and applications to real data.

### Simulations using UK Biobank target samples to assess extension of S-LDSC to low-frequency variants

To assess possible biases in heritability and enrichment estimates under a more realistic scenario, we simulated quantitative phenotypes from chromosome 1 of UK Biobank interim release dataset with imputed variants from 1000G and UK10K (113,851 unrelated individuals, 1,023,655 variants with allele counts greater or equal to 5 in UK10K). First, we randomly sampled integer-valued genotypes from UK Biobank imputation dosage data. Second, we set trait heritability to *h*^2^ = 0.5, selected *M* = 100,000 causal variants, and performed simulations under a coding-enriched architecture by simulating the variance of per-normalized genotype effect sizes proportional to 1_*j non-coding*_(2*p*(1 #x2212; *p*))^1+*α*_0_^ + *c* * 1_*j coding*_(2*p*(1 − *p*))^1+*α*_*coding*_^, where 1_*j coding*_ (resp. 1_*j non-coding*_) indicator function taking the value 1 if variant *j* belongs (resp. does not belong) to the coding annotation, *p* is the frequency of the causal variant in the simulated UK Biobank genotypes dataset, *α*_0_ was set to −0.25, and *c* and *α*_*coding*_ were chosen to produce four different genetic architectures (see Table S4). We note that this generative model is different and more complex than the additive inference model implemented in S-LDSC, but may be more realistic as the effect size of coding variants depends now directly on their allele-frequency (and not or their low-frequency/common status). We also performed simulations under an enhancer-enriched architecture by considering the baseline ChromHMM/Segway weak-enhancer^37^ annotation, which has similar properties as the coding annotation (2.28% of reference low-frequency variants versus 1.83% for coding, and elements with a mean length size of 249bp versus 315bp for coding). To investigate the impact of the LD-dependent architecture created by the enrichment of these two annotations (coding and weak-enhancer variants tend to have low levels of LD^31^), we randomly created 100 shifted coding (resp. weak-enhancer) annotations, and selected the annotation with an average level of LD (i.e. the shifted annotation with the 50^th^ smallest level of LD computed on low-frequency variants; see ref. ^31^ for a definition of level of LD). Third, we used version 2.3 of BOLT-LMM software^24,88^ (see URLs) to compute association statistics on UK Biobank dosage data to mimic the fact that we computed summary statistics on imputed data. Finally, we used S-LDSC with our baseline-LF model (except that the 6 new functional annotations were not included in the simulation analyses) with UK10K as an LD reference panel to estimate 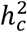, 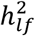, and coding/enhancer CVE and LFVE. S-LDSC was run by restricting regression variants to accurately imputed variants (i.e. INFO score^36^ ≥ 0.99), as we suggested previously^5^, or to all variants (irrespective of INFO score). We also report results when using an INFO score threshold of 0.5 or 0.9 (see Table S4). We also considered including INFO score explicitly in the regression to down-weight poorly imputed variants (i.e. replacing equation (1) by 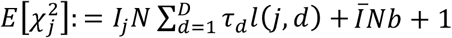, where *I*_*j*_ is the INFO score of variant *j* and 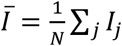; this approximation assumes that genotype uncertainty decreases the association test statistics), but this did not improve the results, consistent with the fact that summary statistics computed from dosage data already down-weight poorly imputed variants (Table S4). Our simulation framework is summarized in Figure S2. We performed 1,000 simulations for each simulation scenario. In each case, we removed 0-3 outlier simulations in which the estimate of 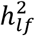 was below 0.0001; we did not observe any such outlier results in analyses of real traits (minimum 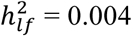 = 0.004; Table S5).

### UK Biobank data set and choice of traits for main analyses and meta-analyses

We analyzed data from the UK Biobank^22,23^ consisting of 487,409 samples genotyped on ~800,000 markers and imputed to ~93 million SNPs using the Haplotype Reference Consortium dataset^25^ (*N* = 64,976 haplotypes from WGS data). We restricted our analyses to 408,963 individuals with UK ancestry^23^ and to variants with MAF ≥ 0.1% that were present in our UK10K LD reference panel (8,523,464 variants in total). Overall, the imputation quality was high even for low-frequency variants (median INFO score larger than 0.9 for variants with MAF between 0.5% and 0.7%; Figure S17). For each investigated trait (see below), we computed mixed model association statistics using BOLT-LMM version 2.3 software^24,88^ with genotyping array (UK BiLEVE / UK Biobank), assessment center, sex, age, and age squared as covariates. We also included 20 principal components to correct for ancestry (provided with the UK Biobank data release^23^) according to guidelines of ref. ^24^. We included 672,292 directly genotyped SNPs in the mixed model (specifically, all autosomal biallelic SNPs with missingness < 10 %).

We restricted our analyses to UK Biobank traits for which the *z*-score for nonzero 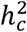 computed using S-LDSC with the baseline-LF model was at least 10, to maximize robustness of 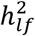 (Table S5). We selected 40 traits (average *N* = 363,166) with squared phenotypic correlation below 0.5 (except for diastolic and systolic blood pressure phenotypes, which are traditionally analyzed jointly; Table S17). In all meta-analyses, we excluded one of each pair of traits with squared phenotypic correlation larger than 0.1, prioritizing traits with larger *z*-score for nonzero 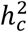 (27 independent traits; average *N* = 354,892). All meta-analyses were performed using random-effects meta-analyses implemented in the R package *rmeta*.

### S-LDSC analyses of UK Biobank data

We applied S-LDSC with the baseline-LF model to 40 UK Biobank traits, estimated 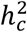, 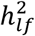 and the 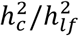 ratio using the 15 MAF bin annotations, and computed their standard errors using a jackknife procedure. We meta-analyzed the 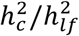 ratio, and multiplied it by the ratio of the number of low-frequency and common variants in the LD reference sample (i.e. 3,398,397/5,353,593) to convert it into a per-variant heritability ratio. To match these ratios to a model in which the variance of per-normalized genotype effect sizes is proportional to (*2p*(1 − *p*))^1+α^, we used low-frequency and common variants of our LD reference panel and computed the 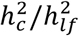 ratio using different values of *α*. We note that BOLT-LMM uses a linear mixed model and achieved an effective sample size (*N*^*eff*^) higher than the true sample size (*N*) (ref. ^24^). This implies that S-LDSC overestimates per-variant heritability when using summary statistics generated by BOLT-LMM. We thus corrected 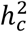 and 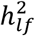 estimations by dividing them by *N*^*eff*^/*N*, with *N*^*eff*^ estimated by taking ratios of chi-square statistics computed by BOLT-LMM vs. linear regression at genome-wide significant SNPs (as in ref. ^24^). After this correction, our reported 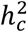 and 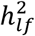 estimates are only 1.15x and 1.16x higher than when applying S-LDSC to summary statistics computed using linear regression with 20 principal component covariates on *N* = 337,539 unrelated individuals with UK ancestry, and our 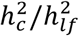 ratio estimates are nearly identical 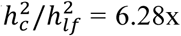, s.e. = 0.23x with BOLT-LMM vs. 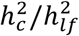 = 6.34x, s.e. = 0.26x with linear regression; Figure S18), confirming that our results are not impacted by the use of BOLT-LMM summary statistics.

The CVE and LFVE of each functional annotation were compared using a *z*-test; these values are independent as they are computed using non-overlapping sets of variants. The regression slope of LFVE on CVE was computed with no intercept. As most of the 33 annotations are correlated, we did not attempt to assess the statistical significance of the regression slope, or of the corresponding correlation between CVE and LFVE.

For CTS analyses, we analyzed the 396 Roadmap^7^ annotations constructed in Finucane et al.^48^ from narrow peaks in six chromatin marks (DNase hypersensitivity, H3K27ac, H3K4me3, H3K4me1, H3K9ac, and H3K36me3) in a subset of a set of 88 primary cell types/tissues. We selected CTS annotations for which common variants are disease relevant following Finucane et al.^48^ guidelines. First, we analyzed each CTS annotation in turn using default S-LDSC (i.e. not our extension to low-frequency variants) by conditioning on all the non-CTS annotations of the baseline-LD model v1.1, the union of annotations for each of the six chromatin marks, and the average of annotations for each mark (as performed in ref. ^48^). We note that our choice to switch from the baseline model^5^, as performed in ref. ^48^, to the baseline-LD model (which includes MAF bins and LD-related annotations in addition to new functional annotations) was motived by our observation that the baseline model can slightly overestimate functional enrichment due to unmodeled annotations^31^. We also decided to consider only non-CTS annotations and to remove the four enhancers annotations derived from Vahedi et al.^89^ (absent from the baseline model and added in the baseline-LD model) as they are T-cell specific and may impact the detection of relevant cell types for traits for which T cells are a relevant cell type (such as asthma and eczema; see Figure S19). We retained all the CTS annotations with a *τ* coefficient statistically larger than 0 (using *P* < 0.05/396), selecting a total of 637 trait-annotation pairs with at least one CTS annotation for 36 of 40 traits (all traits except high light scatter reticulocyte count, high cholesterol, sunburn occasion, and age at menopause), including 25 of 27 independent traits (Table S9). Finally, we re-analyzed these 637 trait-annotation pairs using our extended S-LDSC with the baseline-LF model, the union of the six chromatin marks, and the average of annotations for each mark. In Figure 4, we report all 637 pairs for completeness, demonstrating the consistency between CVE and LFVE for CTS annotations (Table S10). However, as the 1-53 CTS annotations selected for each trait are often highly correlated with each other, we selected for each of the 25 independent traits the “most critical” CTS annotation, defined in the main text and Figure 4 as the CTS annotation with the most statistically significant CVE. For these 25 annotations, we regressed their LFVE on their CVE with no intercept. We also considered 4 alternative definitions of the “most critical” CTS annotation for each trait: 1) the CTS annotation with the highest CVE, 2) the CTS annotation that explains the higher proportion of 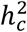, 3) the CTS annotation with the most significant *τ* coefficient, or 4) the CTS annotation with the highest standardized *τ*\* coefficient (defined as the proportionate change in per-variant heritability associated to a one standard deviation increase in the value of the annotation^31^). We also computed 5) the mean LFVE and CVE across 1-53 significant CTS annotations for each independent trait. For each of these definitions, LFVE were similar to CVE (Figure S9). Finally, when testing if a CTS annotation has a significantly larger LFVE than CVE, we used a trait-specific Bonferonni threshold (i.e. 0.05 divided by the number of CTS annotations retained for the trait).

For gene set analyses based on the *s*^*het*^ metric^30^, we divided variants into 5 bins containing the same number of genes (3,073; 3,072 for the last bin). For S-LDSC analyses, we added to the baseline-LF model two annotations for variants inside a protein coding gene (for low-frequency and common variants, respectively; we used the 17,484 protein-genes from ref. ^51^), 10 annotations for variants inside the 5 gene sets, and 10 annotations for non-synonymous variants inside the 5 gene sets (22 annotations in total).

### Forward simulations

To investigate the connection between LFVE, CVE and the distribution of fitness effects (DFE), we performed forward simulations under a Wright-Fisher model with selection using SLiM2 software^59^ (see URLs). We simulated 1Mb regions of genetic length 1cM with a uniform recombination rate and a uniform mutation rate (2.36 × 10^−8^, as recommended in SLiM manual). *De novo* mutations had probability π_*del*_ to be deleterious with a dominance coefficient of 0.5 and a selection coefficient *s* drawn from a gamma distribution with mean 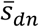 and shape *θ*, and had probability 1 - π_*del*_ to be neutral (i.e. *s* = 0). We outputted a sample of 5,000 European genomes using the out-of-Africa demographic model of Gravel et al.^60^ implemented in SLiM. Then, we used Eyre-Walker model^10^ to compute the per-allele effect size 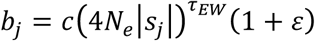, where *c* is a constant, *N*_*e*_ is the effective population size, *s*^*j*^ the selection coefficient of variant *j*, *τ*_*EW*_ is the coupling coefficient between selection and phenotypic effect, and *ε* is a normally distributed noise. Here, *c* was set to have a trait heritability *h*^2^ = 0.5 (i.e. 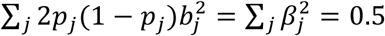, where *p*_*j*_ is the allele frequency of variant *j*), *N*_*e*_ was set as the expected coalescent time^90^ of the European population of the Gravel et al. model (6,524), and *ε* was set to 0 for simplicity. We note that we focused here on per-variant heritability (i.e. 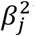) and not directional effects, thus our conclusions are independent of the direction of the selection coefficient on the trait and are valid for traits that are either under direct or stabilizing selection.

Unlike our previous forward simulation framework^31^, we designed these simulations to have a realistic DFE for annotations mimicking both non-synonymous and non-coding variants. We first performed simulations involving non-synonymous and non-coding variants, in order to fit appropriate parameter values for these annotations; subsequent simulations distinguishing functional non-coding and ordinary non-coding variants are described below. We created 50 non-synonymous elements with a realistic length 200bp (10kb in total, 1% of the 1Mb simulated genome) separated by non-coding elements of size 14.9kb (99% of the simulated genome; Figure S11a). To mimic non-synonymous elements, we used π_*del*_ = 80%, 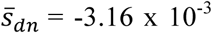 and *θ* = 0.32, as previously estimated^11^. As the DFE of non-coding regions has never been investigated (to our knowledge), we performed simulations using a wide range of non-coding DFE to find the best fit to observed patterns of enrichment. We varied π_*del*_ for non-coding variants between 5% and 80% (16 values in total using 5% incremental steps), 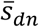 for non-coding variants between - 10^−5^ and −10^−3^ (7 values in total using logarithmic incremental steps), *θ* for non-coding variants between 0.05 and 1 (5 values in total using logarithmic incremental steps), and *τ*_*EW*_ between 0.025 and 1.00 (40 values in total using 0.025 incremental steps), leading to a total of 22,400 different scenarios (Table S18). We simulated 100 regions of 1 Mb for each scenario and merged the outputted variants. We considered all deleterious variants as causal, generated per-allele and per normalized genotype effect size using the Eyre-Walker model, and removed scenarios in which our simulated values of per-variant heritability ratio between common and low-frequency variants, non-synonymous LFVE, and non-synonymous CVE were outside the 95% confidence intervals estimated by the meta-analysis of UK Biobank phenotypes (i.e. 3.99x [3.69; 4.29]), 38.25x [33.80; 42.70] and 7.72x [6.05; 9.39], respectively). After those filters, 11 scenarios remained; π_*del*_ were between 0.30 and 0.75, 10/11 scenarios had 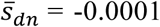, 9/11 scenarios had *θ* = 0.32, and *τ*_*EW*_ between 0.675 and 0.925. We selected the scenario in which the simulated values were closest to the values from the UK Biobank meta-analysis based on Euclidian distance (Table S19); more precisely, we retained the scenario where π_*del*_ = 40%, 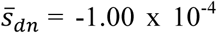, *θ* = 0.32 for non-coding variants and *τ*_*EW*_ = 0.75 and obtained the values 4.07x, 36.27x and 7.73x for per-variant heritability ratio between common and low-frequency variants, non-synonymous LFVE, and non-synonymous CVE, respectively. These estimated parameter values suggest that a *de novo* non-coding mutation is only 2 times less likely to be deleterious than a *de novo* coding mutation, but on average has a selection coefficient 30 times smaller. They also predict a non-synonymous rare variant (MAF < 0.5%) enrichment of 74.27x (Table S19). However, we caution that the diversity patterns in this simulation scenario were not realistic. We observed a ratio between the number of low-frequency and common variants of 0.89, compared to 0.63 in UK10K data (3,398,397 low-frequency variants against by 5,353,593 common variants), and a ratio between the proportion of non-synonymous variants in low-frequency and common variants of 1.38, compared to 1.65 in UK10K data (non-synonymous variants represent 0.45% of low-frequency variants and 0.27% of common variants). Indeed, the Gravel et al. demographic model^60^ that we used is based solely on the site frequency spectrum of non-synonymous variants, which becomes unrealistic when considering other variants. However, we believe that this issue does not impact our conclusions, which are based on the per-variant heritability of an annotation rather than the proportion of heritability explained by the annotation. We also note that our *τ*_*EW*_ estimate (0.75) is much larger than previous estimates^11,14,15,58^, primarily due to the fact that we explicitly model different distributions of selection coefficient for non-synonymous and non-coding variants. To assess the robustness of this estimate, we randomly resampled the values of per-variant heritability ratio between common and low-frequency variants, non-synonymous LFVE, and non-synonymous CVE from their estimated values and standard errors (1,000 resampling sets) and re-estimated *τ*_*EW*_ using a similar procedure. We observed that *τ*_*EW*_ estimates tended to be a slightly larger than 0.75 (Figure S15), confirming the high coupling coefficient between selection and effect sizes.

In most subsequent simulations, we fixed the probability of a deleterious variant to be causal (π_*del:causal*_) at 10%, so that the proportion of *de novo* non-synonymous variants that are causal (π, defined as π = π_*del*_·π_*del:causal*_) is 8% (resp. 4% for non-coding variants). This allows non-synonymous variants to have LFVE and CVE on the same order of magnitude as the LFVE and CVE observed for the non-synonymous variants inside genes predicted to be under strong negative selection^30^ (102.0x and 41.4x, respectively; Figure 5). We note that we replicated our main results when using π_***del:causal***_ = 5% (Figure S20).

Next, we investigated the impact of 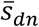 and π on a “functional non-coding” annotation. To do so, we alternately considered 200kb functional elements as non-synonymous elements (1% of the simulated genome) or as functional non-coding elements (1% of the simulated genome), separated by “ordinary non-coding” elements of size 9.8kb (98% of the simulated genome; Figure S11b). For each functional non-coding element, we fixed π_*del*_ = 60% and *θ* = 0.32 (equal to the value of *θ* for non-synonymous and overall non-coding elements). We chose a value π_*del*_ in between the value for overall non-coding (π_*del*_ = 40%) and non-synonymous (π_*del*_ = 80%) annotations, as we hypothesized that enriched functional non-coding annotations in the human genome have a larger proportion of deleterious variants than the overall non-coding genome. However, we note that we obtained similar results when choosing π_*del*_ = 40% for the functional non-coding annotation (Figure S21). We varied 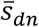 and π_*del:causal*_ (and thus π) of the functional non-coding annotation, while retaining π_*del:causal*_ = 10% for the variants in the non-synonymous and ordinary non-coding elements. (We varied 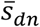 on the logarithmic scale, and report truncated values in the manuscript for simplicity; for example, 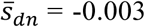 stands for −3.1623 × 10^−3^; see Table S12 for exact 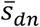 values). For each scenario, we simulated 1,000 regions of 1Mb for each scenario, merged the outputted variants, and considered 100 randomly chosen sets of causal variants.

## URLs

ldsc software, http://www.github.com/bulik/ldsc.

baseline-LF annotations: https://data.broadinstitute.org/alkesgroup/LDSCORE/baselineLF.

BOLT-LMM association statistics computed in this study will be made available for public download via the UK Biobank Data Showcase.

phastCons elements, http://hgdownload.cse.ucsc.edu/goldenPath/hg19/database/phastConsElements46way*.txt.gz;

Flanking bivalent TSS/enhancers, http://egg2.wustl.edu/roadmap/data/byFileType/chromhmmSegmentations/ChmmModels/coreMarks/jointModel/final/*_15_coreMarks_segments.bed.

BOLT-LMM software, https://data.broadinstitute.org/alkesgroup/BOLT-LMM.

SLiM2 software, https://messerlab.org/slim/.

## Acknowledgments

We thank A. Gusev, Y. Reshef, F. Hormozdiari, O. Weissbrod, B. Neale, A. Siepel and S.M. Gazal for helpful discussions. This research has been conducted using the UK Biobank Resource (application number 16549). This research was funded by NIH grants U01 HG009379, R01 MH101244, R01 MH107649, R01 MH109978 and U01 HG009088.

